# Validation of plasmonic-based biosensors for rapid and sensitive detection of rabbit hemorrhagic and foot-and-mouth disease viruses in biological samples

**DOI:** 10.1101/2024.04.15.589493

**Authors:** Chiara Urbinati, Vittoria Di Giovanni, Giulia Pezzoni, Lorenzo Capucci, Marco Rusnati

## Abstract

Biosensing technologies and monoclonal antibodies (MAbs) are gaining increasing importance as powerful tools in the field of virology. Surface plasmon resonance (SPR) is an optical biosensing technology already used in virus detection and in the screening of MAbs of diagnostic and therapeutic value. Rabbit haemorrhagic disease virus 2 (RHDV) and foot-and-mouth disease virus (FMDV) are top veterinary issues for whom, the development of novel methods for their detection in biological samples represents a priority with important livestock healthcare and economic implications. With these premises, here we prepared a series of SPR biosensors containing RHDV2 or its 6S subunit immobilized to the surface by different strategies. The biosensors were then used to characterize the binding capacity of a panel of anti-RHDV2 MAbs. From the comparison of the results obtained, the biosensor composed of intact RHDV2 captured with catcher-MAb covalently immobilized to the surface showed the best analytical performances, that were retained also when the same strategy was adopted to prepare a biosensor containing a different virus (namely, FMVD). The results obtained are discussed in view of the exploitation of SPR in the rapid, sensitive and resilient detection of viruses in biological materials and in the screening of antiviral MAbs libraries.

---

Viruses are the cause of important human and animal diseases worldwide, representing a top global healthcare and economic problem. Also, zoonotic spillovers has come to the limelight as a serious threat to human healthcare, adding further importance to the detection of viruses and viral antigens in biological samples^[1,2].^ Currently, diagnosis of viral diseases and detection of viruses or viral antigens in biological samples are still based on laboratory tests, mainly RT-PCR reaction and enzyme-linked immunosorbent assay (ELISA) reaction. However, an increasing shifting away from these technologies is occurring due the requirement of more powerful and automated tools. Biosensing technologies have emerged as promising for reliable and automated detection of intact viruses or their antigens and for the screening of antiviral monoclonal antibodies (MAbs)^[3,4]^. Biosensors are analytical devices composed of a biological component associated with a substrate that acts as physicochemical detector (optical, piezoelectric, electrochemical) and are used for the detection of compounds ranging from small chemical drugs to nano-objects^[5,6]^.

Among the various biosensing technology, surface plasmon resonance (SPR) is a solid-phase, optical-based, label-free and real-time biosensing technology that allows the evaluation of a wide arrays of interactions of biological and pharmacological interest. It is based on a polarized beam of visible monochromatic light that is passed through a prism fitted with a gold film attached to a glass. When the light hits the glass, an electric field is generated and absorbed by free electrons in the gold film, reducing the intensity of the light that, once reflected by the gold, is detected at the specular (resonance) angle that depends on the refractive index of the material present within 300 nm from the gold film. To evaluate the interaction of two binders, one is immobilized onto the gold film (ligand) and then exposed to the second one injected in the fluidic system (analyte). Their interaction causes a change of the refractive index and of the resonance angle, that is presented as a sensorgram that allows the determination of the stoichiometry of the interaction and the dissociation constant (Kd), inversely proportional to the binding affinity.

SPR has been already employed for detection of viral antigens, natural antiviral antibodies and even intact viruses and in biological samples^[7,9]^ and for the screening of MAbs libraries for diagnostic or therapeutic purposes^[10]^.

Technically, the use of intact viruses in SPR technology deeply influences the “geometry” of the biosensor (namely the choice of the ligand and of its immobilization strategy)^[11]^. Indeed, being viruses complex and fragile structures, they are generally unsuited for harsh procedures of chemical immobilization and biosensor regeneration. Also important, SPR is better performed with analytical-grade reagents, otherwise generating heterogeneous surfaces burdened by aspecific bindings and difficult analysis^[12]^ that limit SPR exploitation for detection of viruses or viral antigens in raw biological samples.

In the field of biosensing, a great effort is already being directed towards the development of biosensors for human infectious diseases, while the animal sector has received less attention, despite the appropriate development of diagnostic tests for animal infectious diseases could deliver significant economic advantages and could help in the prompt detection of zoonotic spillovers, an occurrence whose importance has been recent emphasized by the COVID-19 pandemic.

Based on these considerations, we have here decided to perform our research with rabbit hemorrhagic disease virus (RHDV) and foot-and-mouth disease virus (FMDV).

They are two small round non-enveloped RNA viruses belonging to the Caliciviridae and Picornaviridae families, respectively. RHDV is an emerging virus that infects domestic and wild rabbits causing lethal hepatitis with 90-100% of animals dying within 2-3 days’ post infection^[13]^. FMDV causes a highly contagious disease of cloven-hoofed animals including cattle, pigs and sheep with a significant economic impact^[14]^. For both the diseases, the main preventive action is the careful use of vaccines and the implementation of advanced surveillance systems^[15]^. Also fundamental is their accurate and rapid diagnosis that, so far is mainly based on ELISA and RT-PCR reaction, still lacking dedicated and efficient biosensors.

The aim of this work is to help the development of resilient and versatile SPR biosensors for the detection of RHDV2 and FMDV in raw biological samples and the screening of antiviral MAbs, overcoming the limitations of SPR technology described above.

## METHODS AND MATERIAL

As schematized in figure 1a, the work here presented can be divided into: a “preparative phase” aimed at the preparation of the biological samples containing intact RHDV2 or its artificial subunit (RHDV2-6S) and of the different SPR biosensors. An “analytical phase”, aimed at comparing the analytical potential of the different biosensor at detecting intact RHDV2 or RHDV2-6S in biological samples and at selecting the best biosensor geometry. A “validation phase” in which the SPR-generated results were compared with those obtained by standard ELISA and the selected biosensor geometry was used with another virus (namely FMVD).

**Figure 1.**
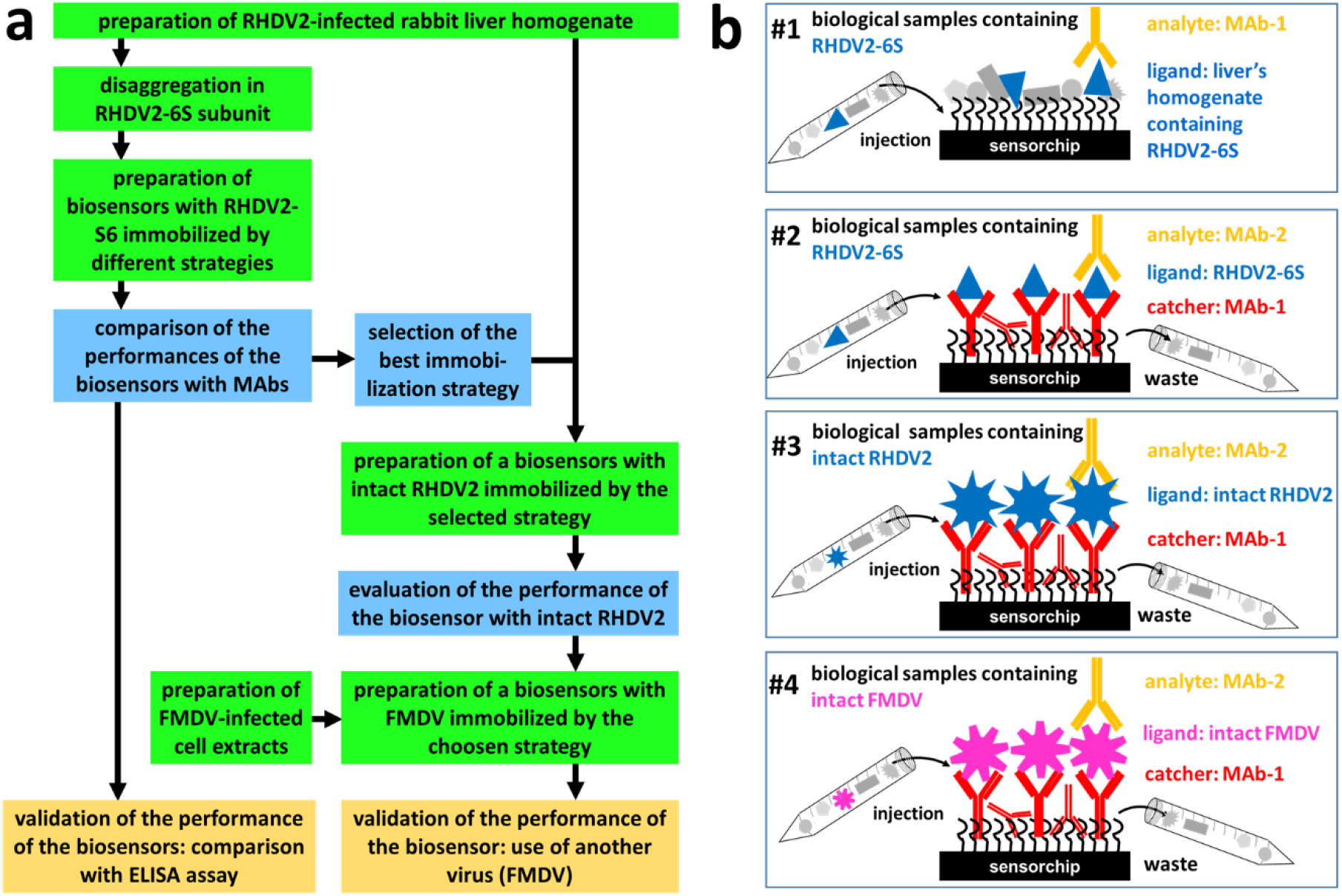
(a) Workflow of the research. Our work is divided into three entwined parts: i) preparation of biological samples and biosensors (green boxes); ii) SPR binding analysis with MAbs to select the best performing biosensor (blue boxes); iii) validation of the selected biosensor by comparing results with those from reference ELISA and by using another virus. (b) Schematic representations of the biosensors used.

### Reagents

Sensorchips CM5 and gel filtration Sephacryl 500HR HiPrep 26/60 column (separation range: 40-20,000 kDa) and Superdex 200 Increase 10/300 GL column (globular protein separation range 10-600 kDa), 1-ethyl-3-(3-diaminopropyl)-carbodiimide hydro-chloride (EDC) and N-hydroxysuccinimide (NHS) were from Cytiva (Marlborough, MA). The panel of mouse MAbs specific for RHDV2 and FMDV were obtained using the method originally described by Milstein and Köhler^[16]^. Anti-RHDV2 MAbs 4H12, 4H7 and anti-FMDV MAbs 1F10, 3B11 and 2A10 are directed against conformational epitopes exposed on the virion, anti-RHDV2 MAb 6G2 is instead directed against a conformational epitope located inside the viral capsid^[17]^. Polyclonal rabbit IgG was purified from the serum of a rabbit convalescent from RHDV2 and possesses a strong protective capacity against RHDV2 due to its high specificity and affinity for viral surface epitopes. The molecular weights (MW) of the reagents used for calculation of molar ratios are: RHDV2-6S: 13.0 kDa; MAbs: 150, kDa; RHDV2: 14,000 kDa; FMDV: 8,000 kDa.

### ELISA

Sandwich ELISA was used to characterize the reactivity of the panel of MAbs against RHDV2, its 6S subunit and FMDV. All ELISA reactions were performed in 96-well plastic plates and had in common: a) adsorption to the plastic surface of catcher rabbit polyclonal IgG or MAb IgG (2.0-3.0 μg/ml) in 100 mM carbonate buffer pH 9.6 by overnight incubation at 4°C; b) three washes with 2x phosphate saline buffer pH 7.4 containing 300 mM NaCl (PBS) and 0.05% Tween-20 for 4 min. at room temperature; c) dilution of the antigens (from 1:33 to 1:72,171) followed by a second MAb conjugated with horseradish peroxidase (HRP), each step in PBS pH 7.4 containing 1.0 % yeast extract and 0.1% Tween-20 with incubation of the plate for 60 min. at 37°C under stirring; d) development of the reaction with 0.5 mg/ml o-phenylenediamine dihydrochloride in citrate buffer pH 5.0 and H_2_O_2_ diluted at 0,02%; e) reading absorbance at 492 nm with a microplate reader (Sunrise™-Tecan, Switzerland).

### Surface plasmon resonance (SPR)

Analyses were performed on a BIAcore X100 instrument (Cytiva) and had in common: a) for amine-coupling immobilization, carboxyl groups of the carboxy methylated dextran matrix of CM5 sensorchips were activated by injecting a 1:1 mixture of 400 mM EDC and 100 mM NHS at 10 μl/min for 7 min. b) after protein immobilization, the excess of reactive groups of the sensorchip were deactivated by injecting 1.0 M ethanolamine at 10 μl/min for 7 min. c) for binding analysis, the “single cycle” procedure was used, in which the harsh regeneration procedure (injection of 10 mM glycine pH 2,5 at 30 μl/min for 30 sec.) is performed only at the end of the complete series of dose-response injection, as to preserve the integrity of the sensorchip^[18]^. Increasing concentrations (0.25, 1.0, 4.0, 20.0 and 100 nM) of the MAbs in 10 mM Hepes, pH 7.4, 150 mM NaCl, 3 mM EDTA, 0,005% surfactant P20 (HBS-EP) were injected at 25°C at 30 μl/min for 120 sec. (biosensor #1 and #2) or for 150 sec. (biosensors #3 and #4) followed by a 600 sec. “dissociation time” in which HBS-EP alone was passed onto the sensorchip. Kd values were calculated by steady state analyses of equilibrium binding by the Langmuir (viz Scatchard) isotherm binding equation performed by the BiaEvaluation program embedded in the BIAcore X-100 instrument which operates by iterative minimizations of chi-square. The biosensors prepared (figure 1b) are described here below:

### Biosensor #1

The sample containing RHDV2-6S was diluted 1:5 in 10 mM sodium acetate, pH 2.8 and injected (85 μl) over flow cell (Fc) #1 of the activated sensorchip at 10 μl/min for 9 min. All the proteins o the sample remain covalently immobilized on the sensorchip in random orientations. On Fc #2, used as negative control and for blank subtraction, the same amount of liver’s homogenate from uninfected rabbits was immobilized in the same experimental conditions.

### Biosensor #2

Mab 6G2 was used as catcher since, recognizing a portion of the inner shell of RHDV2-6S^[17]^, should not interfere with the binding of MAbs recognizing the exposed portion of RHDV2-6S. MAb 6G2 was diluted at 10 μg/ml in 10 mM sodium acetate pH 4.5 and injected (75 μg) over both the Fcs of the activated sensorchip at 10 μl/min for 7 min. The RHDV2-6S-containing sample was injected over Fc #2 of the sensorchip containing MAb-6G2 as described above. In this way, RHDV2-6S remains specifically immobilized onto the biosensor while all the other contaminants pass in the flow through. Fc #1, used as negative control and for blank subtraction was left with only immobilized MAb-6G2.

### Biosensor #3

MAb 4H12, that recognizes an exposed epitope of RHDV2-6S^[17]^, was used as catcher and thus immobilized as described. Due to the multivalence of intact virus, its use as a catcher should not hamper its analysis when used as an analyte. The sample containing intact RHDV2 was injected over Fc #2 of the 4H12-containing sensorchip as described for biosensor #2. The Fc #1, used as negative control and for blank subtraction, was left with only immobilized MAb-6G2.

### Biosensor #4

MAb-2A10 recognizes a neutralizing conformational epitope of FMDV capsid involving VP1 and VP2 structural proteins. It was chosen as catcher MAb and then immobilized as described above. Again, the multivalence of intact FMDV should allow its binding analysis when used as an analyte. The sample containing intact FMDV was injected over Fc #2 of the MAb-2A10-containing sensorchip as described for biosensor #3. Fc #1, used as negative control and for blank subtraction was left with only immobilized MAb-2A10.

## RESULTS

### Preparation of biological samples

The results of the preparation of biological samples can be found in the supporting information (figure S1)

### Evaluation of the reactivity of the MAbs against intact RHDV2 and RHDV2-6s by ELISA

The reactivity of intact RHDV2 with the available MAbs was tested with two sandwich ELISAs differing in the catcher used: rabbit anti-RHDV2 IgG or MAb-6G2 (figure 2a and 2b, respectively). The specificity of the assay is proven by the fact that, independently of the catcher used, unrelated MAb-1H3 does not provide any signal (figure 2a, b). Since anti-RHDV2 rabbit IgGs possesses a strong affinity for viral surface epitopes, the reactivity of MAb-4H12 and 4H7, that recognize epitopes exposed on the virion, is high and identical, while that of MAb-6G2 is 30 times lower, confirming that the epitope is poorly exposed on the virus (figure 2a). When MAb-6G2 is used as a catcher, a general decrease in MAbs-HPR reactivity is observed that is quantitatively different for MAbs-4H12 and -4H7 (3 and 30 times, respectively) (figure 2a, b), thus demonstrating that the epitopes of the two MAbs are distinct and differently exposed on the virus. MAb-6G2 reactivity decreases to almost zero (figure 2b) when acting as both a catcher and HRP tracer.

**Figure 2.**
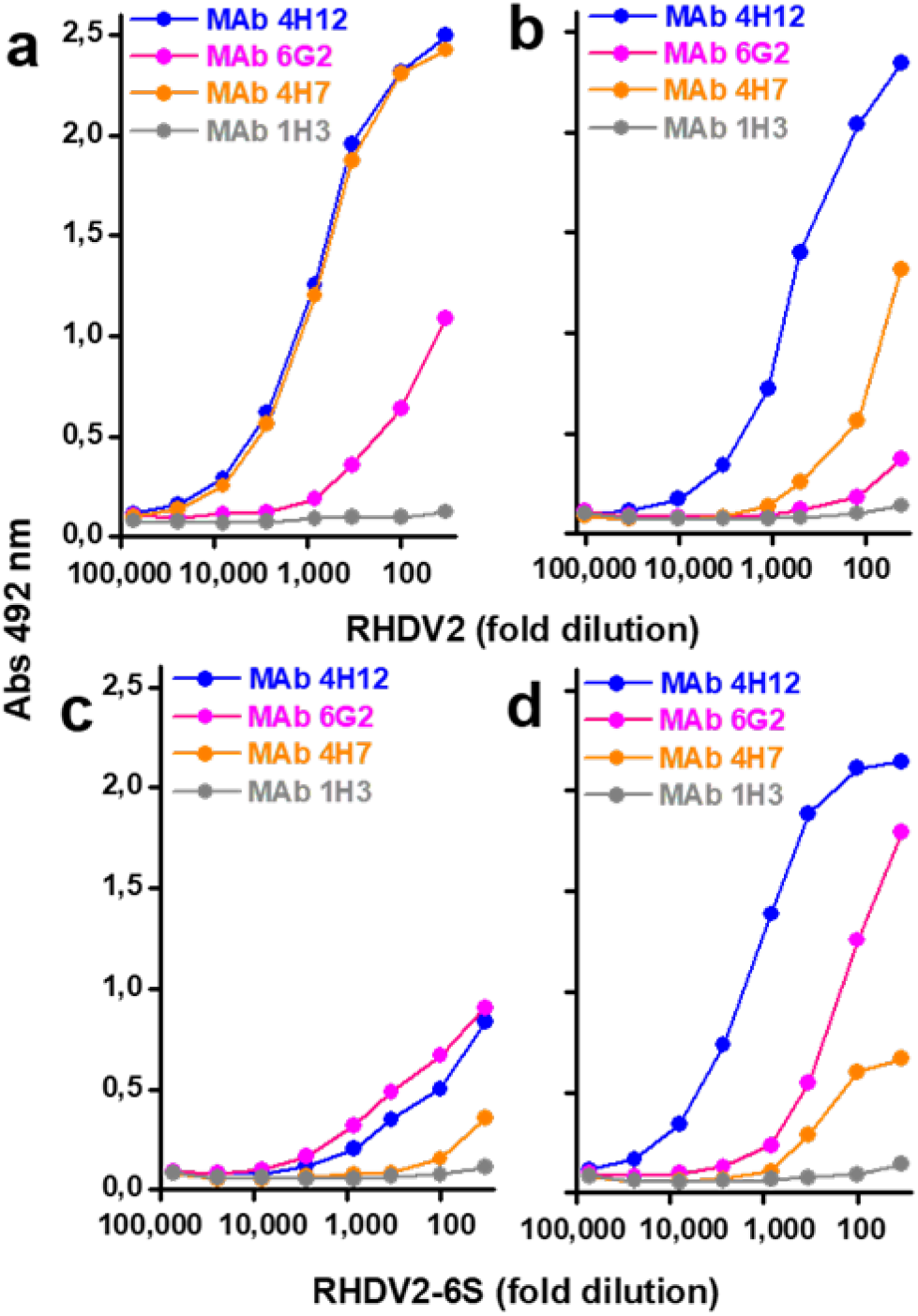
Reactivity of MAbs toward intact RHDV2 and RHDV2-6S subunit by ELISA. HRP-MAb-4H12, 6G2, 4H7 and 1H3 were evaluated for their capacity to bind to RHDV2 (a, b) or to RHDV2-6S (c, d) captured onto anti-RHDV2 IgG (a, c) or MAb-6G2 (b, d).

The reactivity of the MAbs towards RHDV2-6S was tested using the same two ELISA used for the intact virus. Again, the specificity of the assay is proven by the lack of reactivity of unrelated MAb-1H3. Surprisingly, the general reactivity of the MAbs is lower with anti-RHDV2 IgG in respect to Mab-6G2 used as catchers (Fig. 2c, d).

Interesting is the comparison between the two ELISA performed with intact virus and its subunit: with rabbit IgG used as a catcher (Fig. 2a, c), HRP-MAb-4H12 and -4H7 yield a sharp decrease in reactivity for RHDV2-6S in respect to the intact virus (about 50- and 150 times, respectively), confirming the difference between the two epitopes. Mab-6G2 maintains instead a very similar reactivity in the two ELISA (figure 2a, c). With MAb-6G2 used as a catcher, HRP-MAb-4H12 shows a quali-quantitative reactivity 3 times higher for RHDV2-6S in respect to intact virus while HRP-MAb-4H7 shows a 3-times decrease (figure 2b, d), further confirming the epitopes difference. As expected, the reactivity of HRP-MAb-6G2 for the 6S subunit increases (10 times) in respect to intact virus since its epitope, buried inside the virion, becomes instead exposed in the isolated sub unit, despite the steric difficulties that MAbs generally encounter in reacting towards smaller antigens such as the VP60 dimer and the fact that the same MAb is used simultaneously as catcher and tracer.

### Preparation and analytical performances of the biosensors

We firstly prepared biosensor #1, on which liver homogenate containing RHDV2-6S was amine-coupled onto Fc #2, (supporting figure S2a) leading to the immobilization of about 700 RU of proteins (table 1) of which less than 50% should correspond to RHDV2-6S. A similar amount of immobilized proteins from uninfected rabbit liver’s homogenate is obtained on Fc #1.

**Table 1.** Amount of protein/virus immobilized/captured on the biosensors.

| biosensor | immobilized ligand | RU±s.d. | Fmoles/mm<sup>2</sup> | captured ligand | RU±s.d. | Fmoles/mm<sup>2</sup> |
| --- | --- | --- | --- | --- | --- | --- |
| #1 | RHDV2-6S | 666±144 (4) | nd | none | none | none |
| #2 | MAB-6G2 (catcher) | 4,371±2,321 (3) | 29 | RHDV2-6S | 510±397 (17) | 2.8 |
| #3 | MAB-4H12 (catcher) | 3,922±1,183 (5) | 26 | RHDV2 | 3,756±638 (5) | 0.26 |
| #4 | MAB-2A10 (catcher) | 4,514±1,468 (2) | 30 | FMDV | 302±102 (11) | 0.043 |

In brackets: number of immobilizations/captures. nd: not determinable.

The biosensor was then used to assay the available MAbs directed against different epitopes of RHDV2-6S. MAb-4H12 generates a relevant RU increase on both Fcs containing liver’s homogenate from infected or uninfected rabbits (figure 3a). However, the high basal binding does not hamper the determination of a specific binding when the sensorgram is blank-subtracted (figure 3b, c). Indeed, MAb-4H12 binds surface-immobilized RHDV2-6S in a dose-dependent and saturable way with a Kd in the low nanomolar range (table 2). Also the binding of MAb-6G2 is dose-dependent, but occurs with a Kd that is 10 times higher than that of MAb-4H12 (table 2). Both the interactions are specific since unrelated MAb-1H3 binds poorly and in a dose-independent way the biosensor (figure 3c). MAb-4H7 displays an anomalous behavior, following a convex binding isotherm (figure 3c), typical of unspecific low affinity interaction^[19]^.

**Table 2.** Equilibrium binding data for the interaction of the MAbs to the ligands.

| biosensor<br>geometry | MAb | Kd±s.d. (nM) | saturation binding |  |  |
| --- | --- | --- | --- | --- | --- |
|  |  |  | RU max ±s.d. | Fmoles/mm² | MAb:antigen ratio |
| biosensor #1<br>ligand: amine coupled RHDV2-6S<br>analyte: MAbs | 4H12 (4) | 37.0±17.5 | 63.7±28.7 | not determinable |  |
|  | 4H7 (4) | >6.0x106 | 37.5±13.5 | not determinable |  |
|  | 6G2 (3) | 395±209 | 25.7±17.2 | not determinable |  |
|  | 1H3 (2) | no binding |  |  |  |
| biosensor # 2<br>ligand: amine-coupled Mab-6G2,<br>captured RHDV2-6S<br>analyte: MAbs | 4H12 (4) | 15.0±2.3 | 122±31.0 | 0.8 | 1:3 |
|  | 4H7 (5) | 23.3±15.0 | 40.0±29.0 | 0.3 | 1:10 |
|  | 6G2 (2) | no binding |  |  |  |
|  | 1H3 (3) | no binding |  |  |  |
| biosensor #3<br>ligand: amine-coupled MAb-4H12,<br>captured RHDV2<br>analyte: MAbs | 4H12 (3) | 16.2±9.6 | 2,433±603 | 16.2 | 60:1 |
|  | 4H7 (3) | 10.2±6.3 | 1,800±818 | 12.0 | 46:1 |
|  | 6G2 (4) | no binding |  |  |  |
|  | 1H3 (2) | no binding |  |  |  |
| biosensor # 4<br>ligand: amine-coupled MAb-2A10,<br>captured FMDV<br>analyte: MAbs | 1F10 (4) | 19.1±15.6 | 190±101 | 1.3 | 30:1 |
|  | 3B11 (3) | 5.5±4.4 | 265±42.0 | 1.8 | 39:1 |
|  | 2A10 (3) | 8.6±10.3 | 12.0±2.0 | 0.08 | 1:2 |
|  | 2A11 (2) | no binding |  |  |  |

In bracket: number of repeated analysis for each MAB. MAB:antigen ratio is calculated considering RHDV2-6S, intact RHDV2 and FMDV as immobilized ligands for biosensor #1, 2, 3 and 4 respectively.

**Figure 3.**
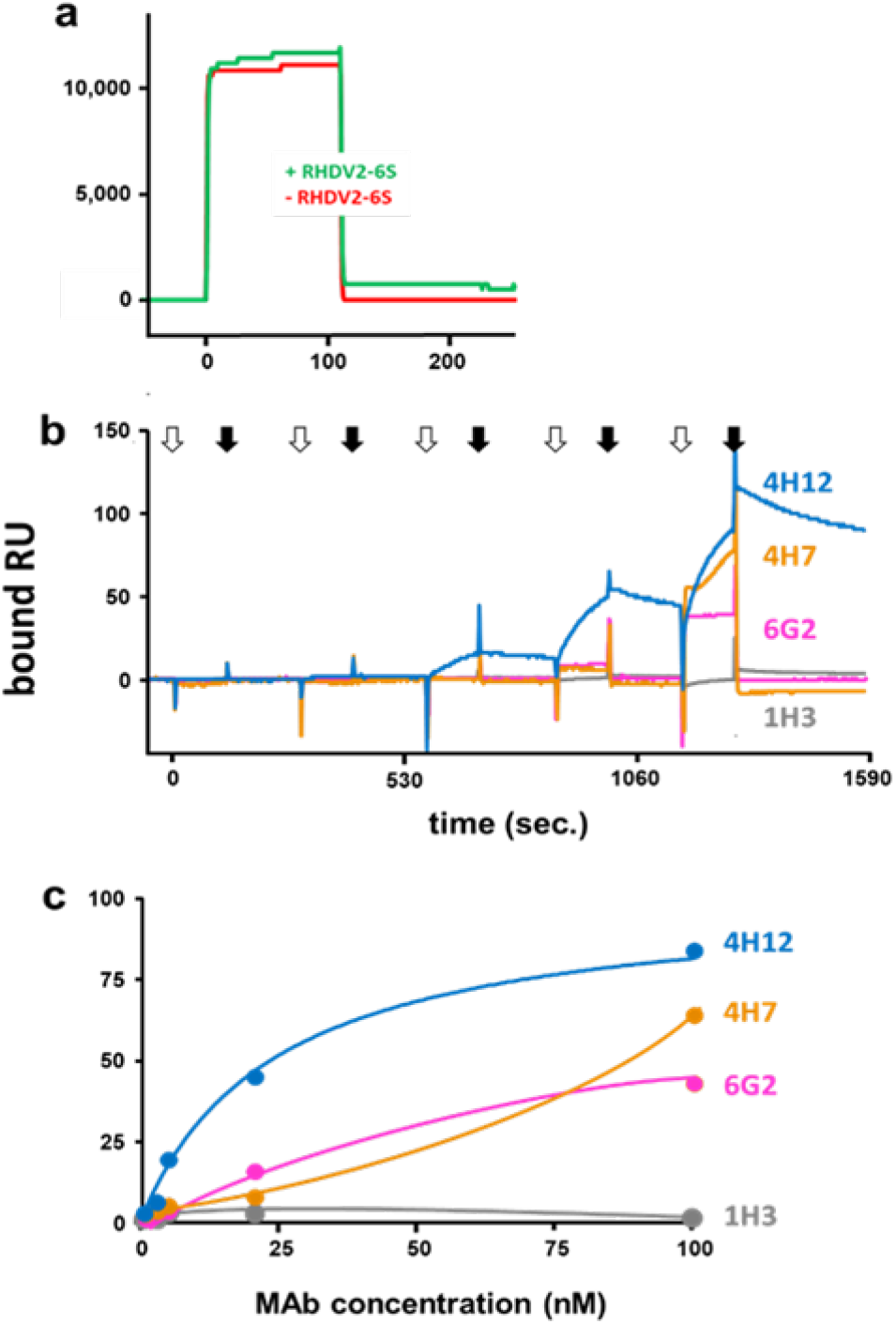
Analytical performances of biosensor #1. (a) Representative sensorgrams showing the binding of MAb-4H12 (100 μM) to liver homogenate from infected (+RHDV2-6S) or uninfected rabbits (-RHDV2-6S). (b) Blank-subtracted sensorgrams of a representative single cycle analysis of the indicated MAbs injected at increasing concentrations on liver homogenate containing RHDV2-6S. White and black arrows indicate start and end of injections, respectively. (c) Steady-state analysis from equilibrium data of panel b.

Biosensor #2 was then prepared by adopting an alternative geometry consisting in the amine coupling immobilization of MAb-6G2 followed by antibody-capture of RHDV2-6S (supporting figure S2b), thus with its exposition in a properly oriented way, possibly mimicking that of the native virion surface. About 4,000 RU (29 Fmoles/mm^2^) of MAb-6G2 remain immobilized that allow the capture of about 500 RU (2,8 Fmoles/mm^2^) of RHDV2-6S (table 1), likely due to the fact that, after random amine coupling immobilization, not all the MAb remains available for antigen binding.

Coming to binding analysis, MAb-4H12 binds surface-immobilized RHDV2-6S in a dose-dependent and saturable way (figure 4) with a Kd comparable to that obtained with biosensor #1 (table 2). Interestingly, with this biosensor, also MAb-4H7 shows a dose-dependent and saturable binding that occurs with a Kd comparable to that of MAb-4H12 (table 2). Instead, MAb-6G2 shows no binding, according to the fact that its epitope is already engaged by the same MAb used as a “catcher”. These bindings are specific since unrelated MAb-1H3 binds poorly to the biosensor.

**Figure 4.**
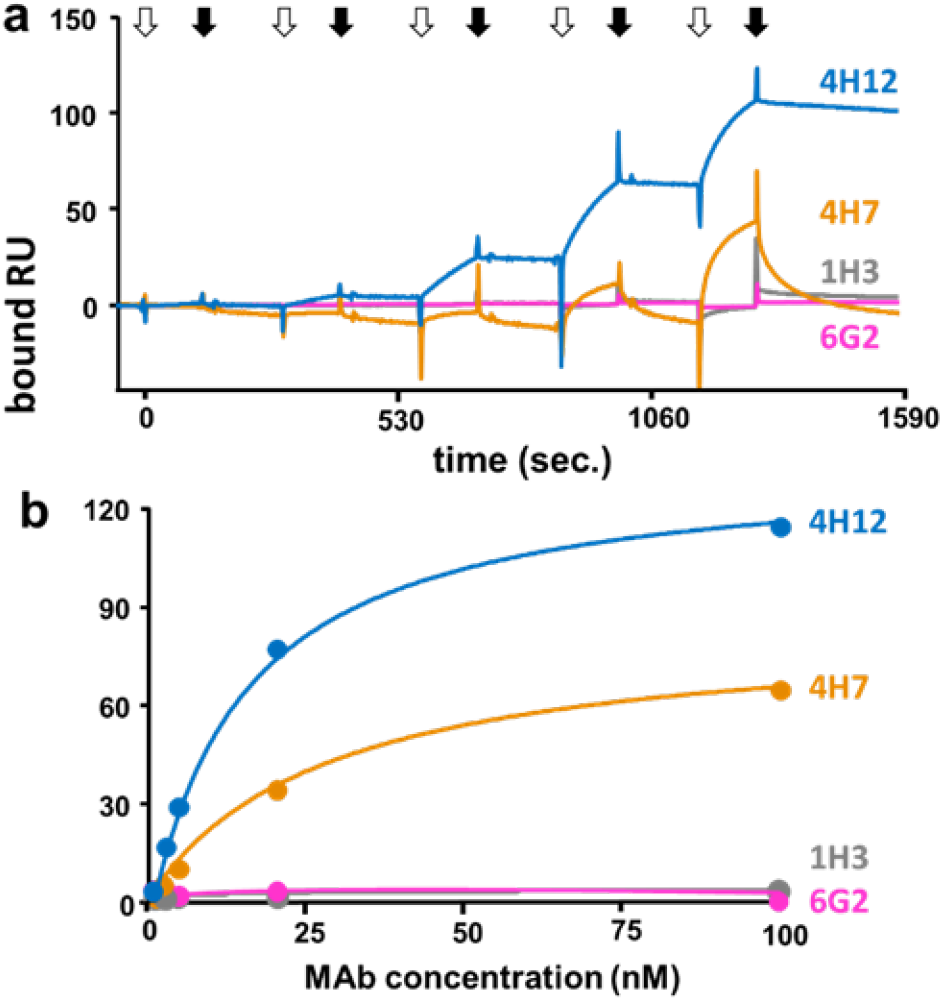
Analytical performances of biosensor #2. (a) Blank-subtracted sensorgrams of a representative single cycle analysis of the indicated MAbs injected at increasing concentrations on RHDV2-6S captured with MAb-6G2. White and black arrows indicate the start and the end of injections, respectively. (b) Steady-state analysis from equilibrium data from panel a.

In conclusion, comparing the analytical performances of the two biosensors, a series of considerations (detailed in discussion) point to biosensor #2 as better suited to identify viral antigens in biological samples and to study MAb/antigen interactions in heterogeneous biological samples, prompting us to evaluate if its geometry was also suited for the identification of intact viruses in biological materials and for the characterization of MAb/virus interaction. To this aim, biosensor #3 was prepared by adopting the MAb-capture strategy with the intact RHDV2. In more details, since MAb-6G2 could not be used as a catcher, being it directed to an epitope that is buried in the intact virus, MAb-4H12 was instead used, grounded on the assumption that virus’ multivalence would ensure the analysis of a MAb also when used simultaneously as ligand and analyte. The amine coupling procedure (supporting figure S2c) leads to the immobilization of an amount of MAb-4H12 that is similar to that obtained for MAb-6G2 (table 1) and that allows a significant capture of RHDV2, as evidenced by the high value of bound RU (table 1). MAb-4H12 and 4H7 bind surface-immobilized RHDV2 in a dose-dependent, saturable way (Figure 5), with a Kd comparable to that obtained for their binding with the isolated RHDV2-6S antigen (table 2). The observed bindings are specific since unrelated MAb-1H3 binds poorly and in a dose-independent way to the biosensor (figure 5). As expected, MAb-6G2 does not bind surface-immobilized RHDV2 since its epitope is buried in the intact virus, being thus unavailable for MAb binding (figure 5). The value of RU bound at saturation for MAb-4H12 and 4H7 to biosensor #3 are 20-40 times higher than those obtained on biosensor #2 and the MAb/RHDV2 ratios are equal to 60:1 and 46:1 for MAb-4H12 and 4H7 respectively (table 2), confirming the expected multivalence of the virus.

**Figure 5.**
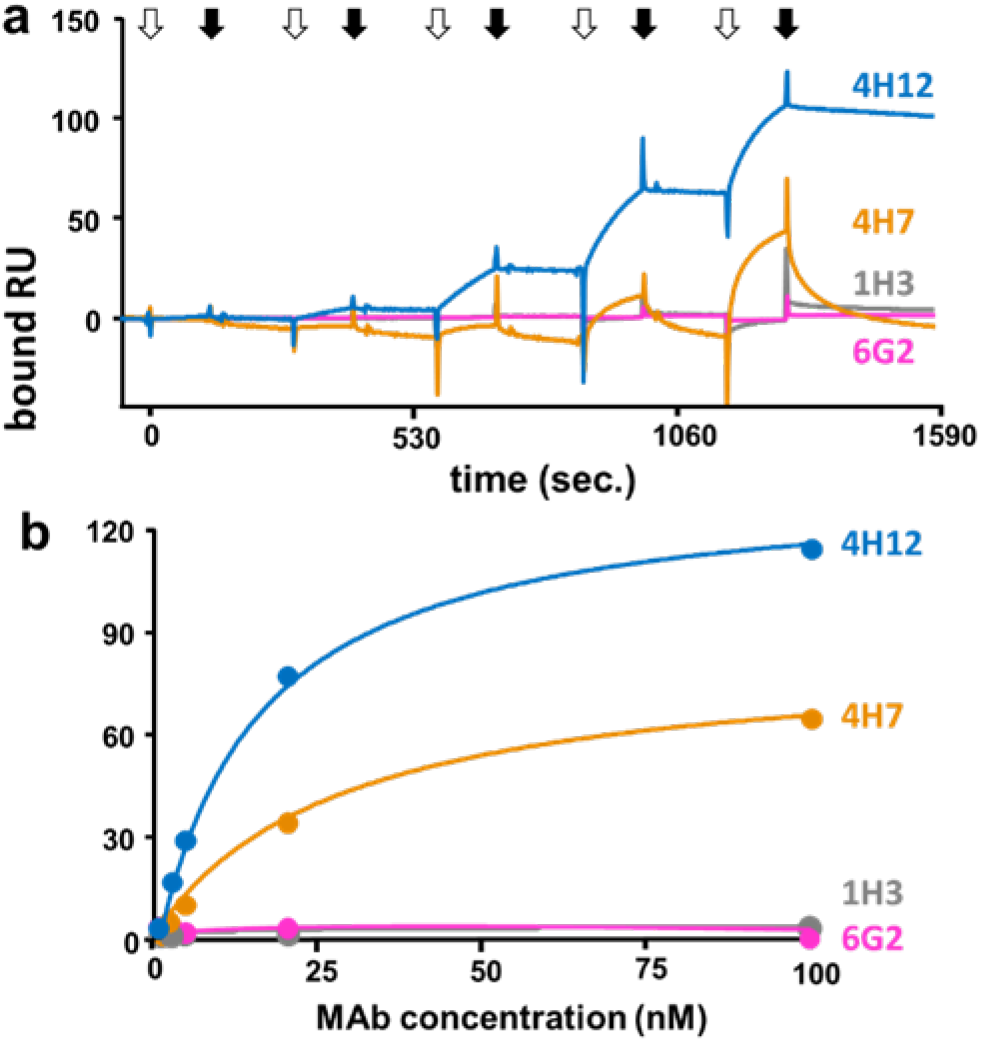
Analytical performances of biosensor #2. a) Blank-subtracted sensorgrams from a representative single cycle analysis of the indicated MAbs injected at increasing concentrations on the surface containing RHDV2-6S captured with MAb-6G2. White and black arrows indicate the start and the end of injections, respectively. b) Steady-state analysis from equilibrium data of panel a.

We then decided to verify if these analytical performances could be replicated with other viruses, as to gain a general validation of the biosensor geometry. To this aim, biosensor #4 was prepared with FMDV. MAb-2A10 was amine-coupled to the sensorchip (supporting figure S2d), leading to an immobilization that is quantitatively comparable to those obtained with other MAbs (table 1). The capture of the FMDV yielded instead a 10 times lower immobilization in respect to that of RHDV2 (table 1).

The biosensor was then challenged with MAbs directed against different epitopes of the FMDV capsid. MAb-1F10, 3B11 and 2A10 bind surface-immobilized FMDV in a dose-dependent, saturable way (figure 6), with comparable Kd values (table 2). The bindings are specific since unrelated MAb-2A11 does not bind to the biosensor (figure 6a, b). Despite a similar binding affinity, the binding capacity (measured as maximal RU bound) of MAb-1F10 and 3B11 is 20-30 times higher than that of MAb-2A10 (table 2). This can be due to a scarce accessibility of MAb-2A10 epitope and could also explain its lower FMDV-capture efficiency.

**Figure 6.**
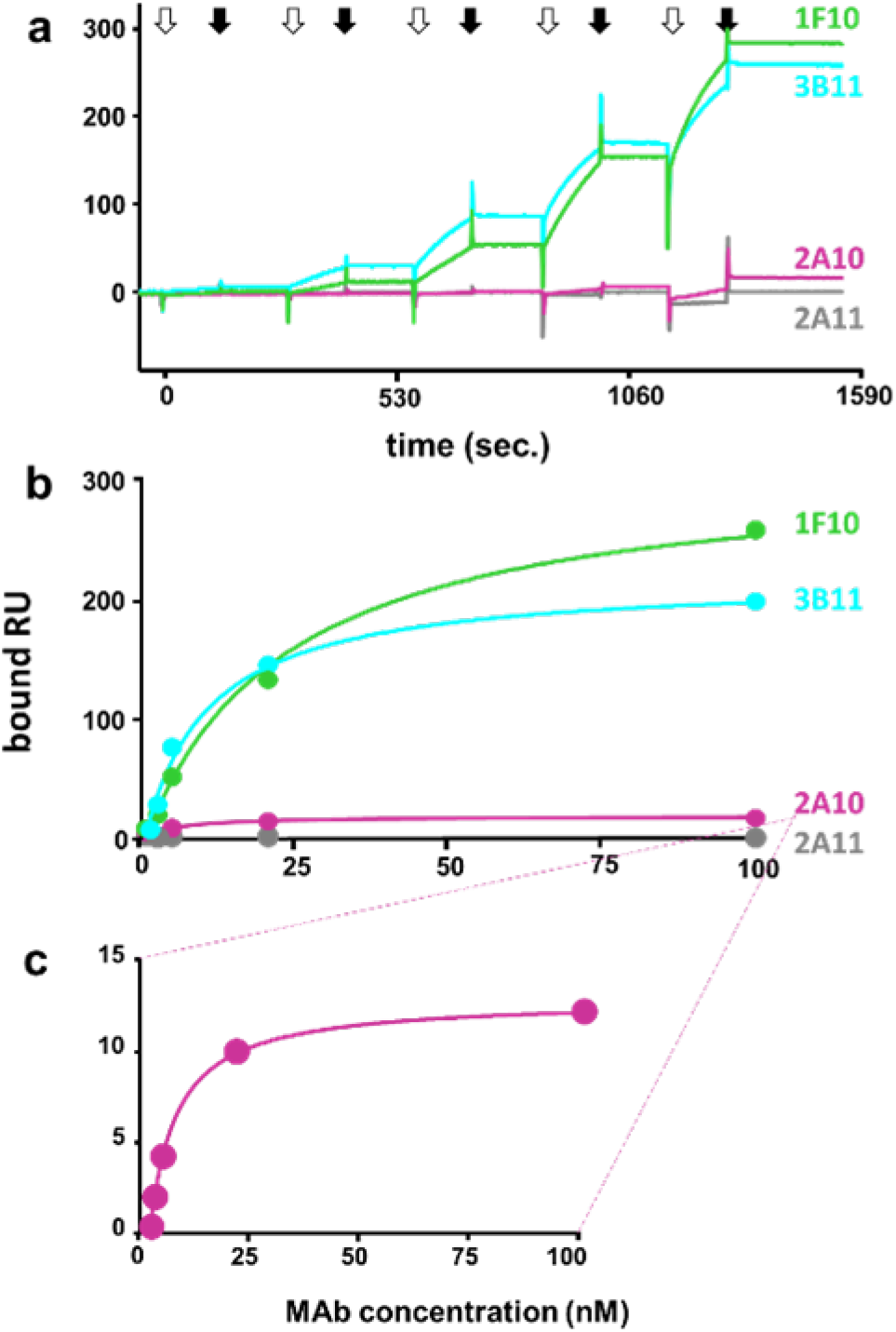
Validation with FMDV. (a) Blank-subtracted sensorgrams of a representative single cycle analysis of the indicated MAbs injected at increasing concentrations on intact FMDV captured with MAb-2A10. White and black arrows indicate the start the and the end of injections, respectively. (b) Steady-state analysis from equilibrium data of panel a. (c) MAb-2A10 has been re-scaled to highlight its dose-dependent, saturable nature.

## DISCUSSION AND CONCLUSIONS

To date, ELISA remains the reference immunoassays for detecting viral antigens, viruses, and related antibodies, owing to its accessibility, low operating and maintenance costs, high sensitivity and specificity. However, ELISA cannot provide direct binding kinetics or affinity values, parameters that are gaining importance with the increasing complexity of the scientific questions. Moreover, ELISA is lengthy, needs the labelling of one binder and appears to be influenced by sera and blood^[20]^. Similarly, also SPR is better suited for analysis of purified binders, limiting its potential in detecting viruses and viral antigens in biological samples. At variance with ELISA, that is performed with disposable materials, SPR biosensors can be used repeatedly, saving reagents and decreasing the cost of the analysis. Unfortunately, however, a repeated use also implies surface-immobilization and regeneration procedures that could be unsuited for fragile structure such as viruses. In this work, we have approached these issues in a comprehensive manner, by preparing and comparing different biosensors for their analytical potential and technical limits at detecting viral antigens or intact viruses in biological samples and at screening panels of MAbs. In the first biosensor prepared, the RHDV2 6S subunit was immobilized to the sensorchip by amine coupling, the most handly and widely used procedure for ligand immobilization[21]. Dealing with a raw liver homogenate as starting material, a heterogeneous surface was obtained that hampered the determination of the stoichiometry of the antibody/antigen interaction. On the bright side, since the calculation of Kd by SPR is independent of the amount of the immobilized ligand, for two out of three MAbs analyzed (4H12 and 6G2), we obtained a saturable, specific binding and related Kd values while for the third MAb, namely 4H7, a non-saturable, aspecific binding was obtained, possibly due to cross-reactivity with other liver proteins present onto the heterogeneous surface of sensorchip. In conclusion, although quick and simple, the direct amine coupling immobilization of a viral antigen contained in a raw biological sample generates a biosensor with significant analytical limits. This prompted us to prepare a second biosensor in which the amine coupling-immobilization of MAb-6G2 allowed the specific capture of RHDV2-6S with the simultaneous riddance of any other contaminant. With this new biosensor a saturable specific binding was obtained also for MAb-4H7 whose Kd was comparable to that of MAb-4H12. In addition, the possibility to determine the exact amounts of captured antigen and bound MAb made possible the calculation of the MAb/antigen ratios, that resulted equal to 1:3 and 1:10 for MAb-4H12 and 4H7 (table 2). This difference could be tentatively due to the close proximity of the 4H7 epitope to that of 6G2, here used as catcher, with its consequent low accessibility. The only limit of biosensor #2 could be found in the fact that when a MAb is used as a catcher (in our case 6G2), it cannot be used also as an analyte, being its epitope on the antigen already engaged. However, it could be anticipated that this would be of no consequence when dealing with multivalent intact virions instead of isolated viral antigens (see below).

Based on these considerations, we decided to prepare a biosensor containing intact RHDV2. Relevant to this aim, the use of amine coupling for intact viruses is burdened by the requirement of extreme pH for efficient ligand immobilization, which can potentially disassemble virions. Also, biosensors containing covalently immobilized ligands usually need harsh regeneration procedures after each analysis, that are scarcely suitable for fragile virion structure. Finally, to avoid heterogeneous surfaces that could make difficult the interpretation of the binding data[12], amine coupling is usually performed with purified virion preparations [22-24]. We then decided to adopt the ligand-capture approach, anchoring the intact RHDV2 virions to MAb-4H12 covalently immobilized to the sensorchip. With this strategy we obtained a biosensor that can be used for a dual purpose: i) the detection of RHDV2 in raw material with significant sensibility, as demonstrated by the high specific signal generated by the injection of RHDV2-infected liver homogenate on biosensor #3 (table 1); ii) the reproducible, long-lasting and meaningful analysis of antiviral MAbs (discussed below). This is allowed by the stability of the covalent immobilization of the catcher MAb and the possibility to replenish the surface with new virus by repeated injections of RHDV2-infected liver homogenate when the analytical performance decreases.

As already mentioned, ELISA still remains the standard assay for virus detection, prompting us to compare our SPR-generated results with those from ELISA, a well-trodden approach already used in many biosensor-oriented works[25]. For what concerns intact RHDV2 analysis, a complete consensus emerges between the two set of results from ELISA and biosensor #3, with negative results for MAb-6G2 and 1H3 and positive results for MAb-4H12 and 4H7, with MAb-4H12 generating higher signals than MAb-4H7. The consensus is retained also for the results for the analysis of RHDV2-6S subunit with ELISA and biosensor #2, with the only exception represented by MAb-6G2, that behaves differently in the two assay: in ELISA, that is performed in static conditions, tracer MAb-6G2 retains a certain capacity to bind RHDV2-6S also when captured by the same MAb, likely due to the tendency of the viral subunit to dimerize[26]. The inability of MAb-6G2 to recognize RHDV2-6s in SPR could be instead tentatively explained by the fact that this analysis occurs under flow, possibly causing a continuous dimer dissociation that hampers the formation of the multimeric binding.

In respect to ELISA, the analysis preformed with the biosensors allowed the real-time calculation of the affinity and molar ratios of the MAb/virus interactions studied that further proved their specificity and effectiveness: i) the affinities of the binding of the two anti-RHDV2-6S MAb-4H7 and 4H12 showed similar for their epitopes when assayed on both biosensor #2 and #3 suggesting their correct accessibility in the two different settings; ii) MAb-6G2, that recognizes an epitope buried inside the virus capsid, binds to biosensor #2 but not to biosensor #3; ii) the amount of MAbs bound at saturation to the biosensor containing the intact virus is 20-40 times higher than that bound to the biosensor containing the isolated viral subunit. This difference is likely due to the possibility that, on the intact virus, the number of exposed RHDV2-6S units is higher than those that is possible to chemically immobilize onto the biosensor. Also, the rounded shape of the virions could increase the binding surface, increasing the number the binding sites available to the MAbs.

In conclusion, we have here developed two biosensors based on antibody capture that demonstrated high potential in the detection of intact virus or viral antigen in raw biological materials and in the screening of MAbs panels that could help in the quick and efficient identification of RHDV2, FMDV and related antibodies in animal population. Also, the exploitation of the biosensor geometry here adopted could be applied to other viruses as well, expanding their use to other areas, also in the human field.

## ASSOCIATED CONTENT

Supporting Information is available free of charge at https://xxxxxxxxxxx

Methods and Material, results, including the chromatographic elution profiles of the preparation of the biological samples used in the work (figure S1) and sensorgrams of the preparation of the biosensors used in the work (figure S2).

## AUTHOR INFORMATION

### Corresponding Author

Marco Rusnati – Department of Molecular and Translational Medicine, University of Brescia, Viale Europa 11 25123, Brescia, Italy; Consorzio Interuniversitario Biotecnologie (CIB), Unit of Brescia, 25123 Brescia, Italy. ORCID: 0000-0001-996878265908;

### Authors

Chiara Urbinati – Department of Molecular and Translational Medicine, University of Brescia, Viale Europa 11 25123, Brescia, Italy.

Vittoria Di Giovanni – Department of Virology, Istituto Zooprofilattico Sperimentale della Lombardia e dell’Emilia Romagna, via Antonio Bianchi 7/9 25124, Brescia, Italy.

Giulia Pezzoni – Department of Virology, Istituto Zooprofilattico Sperimentale della Lombardia e dell’Emilia Romagna, via Antonio Bianchi 7/9 25124, Brescia, Italy.

Lorenzo Capucci – Department of Virology, Istituto Zooprofilattico Sperimentale della Lombardia e dell’Emilia Romagna, via Antonio Bianchi 7/9 25124, Brescia, Italy. ORCID: 0000-0002-1830-3929

Marco Rusnati – Department of Molecular and Translational Medicine, University of Brescia, Viale Europa 11 25123, Brescia, Italy; Consorzio Interuniversitario Biotecnologie (CIB), Unit of Brescia, 25123 Brescia, Italy. ORCID: 0000-0001-9968-78265908;

### Author Contributions

M.R.: writing original draft.; L.C., M.R. supervision, manuscript review and editing, funding acquisition. G.P.: supply of FMDV samples and critical reading of the article. C.U.: methodology and SPR analysis. V.D.G., preparation of RHDV2 biological samples and ELISA testing.

### Notes

The authors declare no competing financial interest.

## ACKNOWLEDGMENTS

This research was founded by Ministero della Salute, Ricerca Corrente, progetto IZSLER 04/18-PRC2018004 to L.C. and M.R. and by Ministero dell’Istruzione, Università e Ricerca (MIUR) (project ex 60%) to MR.

## SUPPORTING INFORMATION

### METHODS AND MATERIAL

#### Preparation of rabbit liver homogenate containing intact RHDV2

Livers from RHDV2-infected rabbits were diluted into 2X PBS at a final liver percentage of 10% w/v, minced with scissors and homogenized with ultraturrax at 4° C, filtered through a gauze, passed through a potter to increase cell breakdown and virus release and centrifuged at 10,000xg for 5 min. at 4°C. The supernatant was collected, centrifuged again at 15,000xg for 30 min. and filtered using a suggestively 0.45 μm cut-off filer paper. RHDV2 positive homogenates were inactivated with 0.8% formalin for 4 h at room temperature and overnight at 4°C. To obtain a sample containing only intact virions, antigenic viral subunits were removed from the raw homogenate by gel filtration chromatography with the chromatographic apparatus Acta Purifier and a Sephacryl 500HR HiPrep 26/60 column (Cytiva, Marlborough, MA, USA) equilibrated with PBS 2x. 30 ml of liver homogenate was loaded onto the column at 2 ml/min. Protein elution was followed by measuring absorbance at 280 nM. 15 ml-fractions were collected and analyzed by ELISA (see below): MAb 4H12 (whose epitope is exposed on the virion) was used to follow the elution of intact RHDV2 MAb 6G2 (whose epitope is buried within the viral capsid) to follow elution of single viral components.

#### Preparation of semi-purified RHDV2 and its RHDV2-6S subunit

Thirty ml of the viral preparation described above was concentrated to 2 ml using a 30 kDa cut-off Millipore filter (Merck KGaA, Darmstadt, Germany) and dialyzed overnight at 4°C in 100 mM carbonate buffer at pH 10. Under these conditions, the RHDV2 viral capsid disintegrates into 6S subunits consisting of 2 VP60 and structurally similar to the native 6S subunit. This was followed by dialysis in PBS 2x for 5 h at 4°C under stirring. A 0,5 ml aliquot of the sample was then loaded onto a Superdex 200 column (Cytiva) equilibrated with PBS 2x at a flow rate of 0,5 ml/min. Protein elution was followed by measuring absorbance at 280 nM. 0.5-ml fractions were collected and analyzed in ELISA as described below.

#### Preparation of FMDV-containing cell extract

The FMDV serotype O strain O1 Manisa was used. Viral antigen was produced starting from baby hamster kidney (BHK)-21 cell line. Briefly, cell cultures at 80% of confluence were inoculated with a viral suspension of about 0.01 multiplicity of infection. After 24-48 hours of incubation at 37°C, when the cytopathic effect was appreciable throughout the entire monolayer, cells were harvested after a freeze-thaw cycle and centrifuged at 5,000xg for 30 min. at 4°C. The supernatant was filtered with a 0.45 μm filter and inactivated by addition of ethyleneimine in form of Binary ethyleneimine as previously described^1^. All the procedures were carried out in a biosecurity level 3 plus laboratory. The effective inactivation of the virus was evaluated according with the “Minimum Bio-risk Management Standards for laboratories working with FMDV” https://www.fao.org/3/cc8479en/cc8479en.pdf).

A 2-liter preparation of FMDV was spiked by an overnight incubation at 4°C with 2% NaCl and 8% PEG 8000. Then, the solution was centrifuged at 8,000xg for 15 min at 4°C and the virus-containing pellet was dissolved in 30 ml of PBS at pH 7.4. The FMDV was then loaded onto a Sephacryl 500HR HiPrep 26/60 column equilibrated with PBS2x at a flow rate of 2,0 ml/min. Protein elution was followed by measuring absorbance at 280 nM. 15,0-ml fractions were collected and analyzed in ELISA as described below.

### RESULTS

#### Biological samples

Liver homogenate from RHDV2 infected rabbits was fractionated on a Sephacryl 500HR HiPrep 26/60 column, resulting in the elution profiles shown in supporting information (figure S1a). As expected from the range of separation of the column, the bulk of proteins elutes in a late peak (fractions 12-16) along with a main peak of viral proteins corresponding to RHDV2-6S, being identified by MAb-6G2. An early immunoreactivity peak also elutes in fractions 7-11 that correspond to intact RHDV2, being identified by its MW (14,500 kDa) and by ELISA with MAb-4H12. This latter peak was pooled and the virus concentration was estimated as 8±4 μg/ml by ELISA titration. Being the total protein concentration equal to 150±25 μg/ml, the degree of virus purification is equal to about 5%.

The virus preparation was dialyzed against a buffer at pH 10.0, that causes virus desegregation in subunits and then fractionated on a Superdex 200 Increase 10/300 GL column. As expected from the range of separation of the column used, the bulk of proteins elutes in the void volume (fractions 2-6), that includes possible residual intact virus. ELISA with MAb-6G2 as catcher and MAb-4H12-HRP as tracer identified an immunoreactivity peak (fractions 7-9, corresponding to proteins with a MW between 100 and 150 kDa), likely consistent with a VP60 dimer. In contrast, this subunit is not detected at all by ELISA when rabbit anti-RHDV2 IgG is used as catcher and MAb-4H7-HRP as tracer, a reaction that is highly positive towards the intact virus (supporting information, figure S1b). RHDV2-6S concentration was estimated was as 50 μg/ml by ELISA tritation. Being total proteins concentration equal to 100 μg/ml, the degree of purity of the RHDV2-6S subunit is equal to 50%.

FMDV infected cell extract was fractionated on a Sephacryl 500HR HiPrep 26/60 column, resulting in the elution profiles shown in supporting information (figure S1c). The absorbance at 280 nm shows that the bulk of proteins elutes in fractions 12-15. Two immunoreactivity peaks were observed: the late one (fractions 13-16), likely corresponding to low MW virus subunits, was discharged, while the early one (fractions 10-12), consistent with intact virus based on its high MW and ELISA results, was used for further analyses. Virus concentration was estimated as 73,0 μg/ml by ELISA titration. Being the total protein concentration equal to 500 μg/ml, the degree of virus purification is equal to about 1,0-2,0 %.

(1) Bahnemann, H.G. Inactivation of viral antigens for vaccine preparation with particular reference to the application of binary ethylenimine. Vaccine 1990, 8, 299-303, doi:10.1016/0264-410x(90)90083-x.

**Figure S1.**
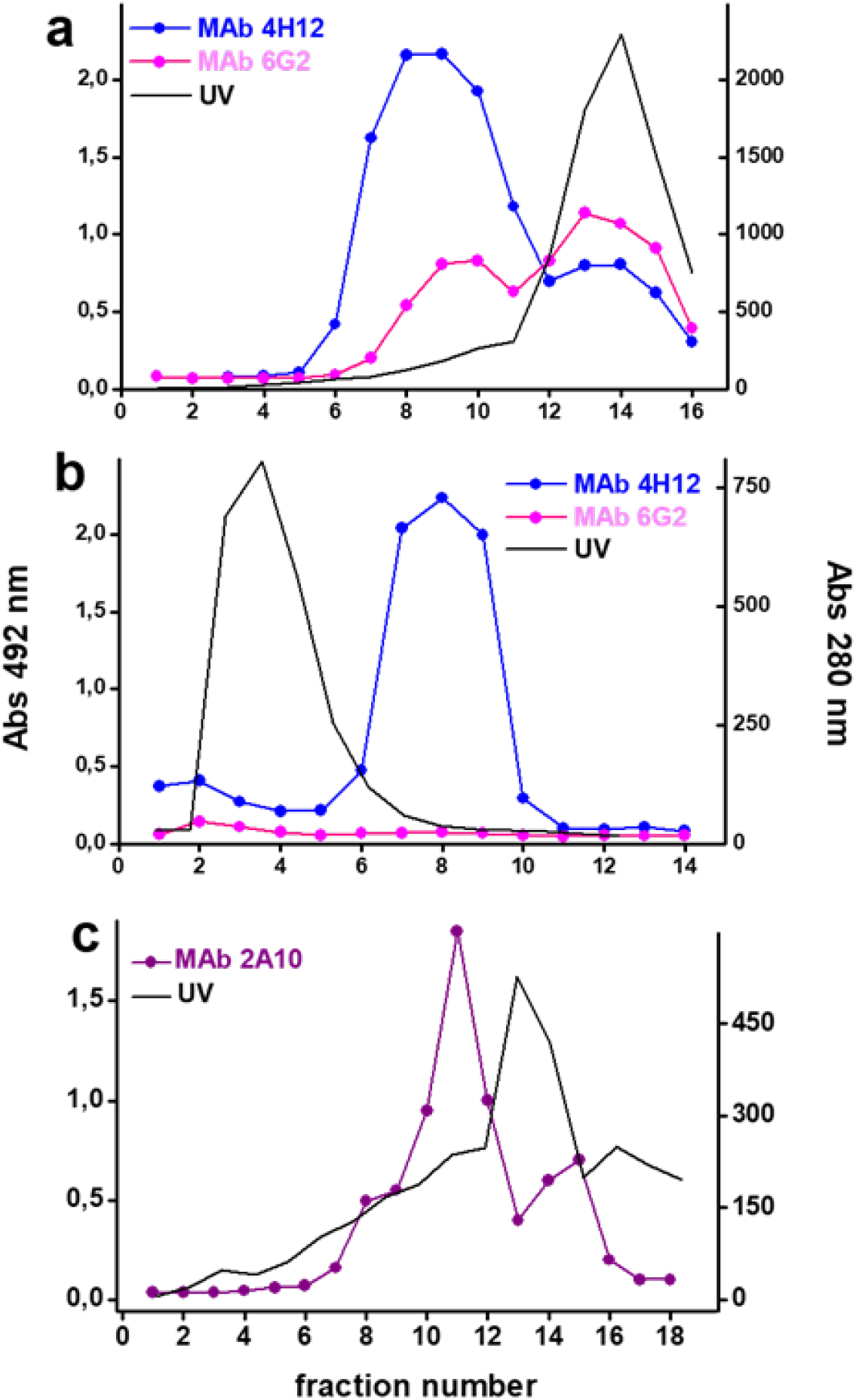
Preparation of biological samples. (a) Intact RHDV2. Liver homogenate from RHDV2-infected rabbits was subjected to gel filtration chromatography monitored by measuring absorbance at 280 nM and by ELISA with catcher-IgG anti-RHDV2 and MAb-4H12-HRP or 6H2-HRP. (b) RHDV2-6S. RHDV2 was disaggregated in RHDV2-6S subunits and subjected to gel filtration chromatography monitored by measuring absorbance at 280 nM and ELISA with catcher-MAb-6G2 and MAb-4H12-HRP or with catcher-IgG anti-RHDV2 and MAb-4H7-HRP. (c) Intact FMDV. FMDV-infected cells extract was subjected to gel filtration chromatography monitored by measuring absorbance at 280 nM and ELISA with rabbit polyclonal antibodies against FMDV type O as catcher and HRP-MAb-2A10 as tracer.

**Figure S2.**
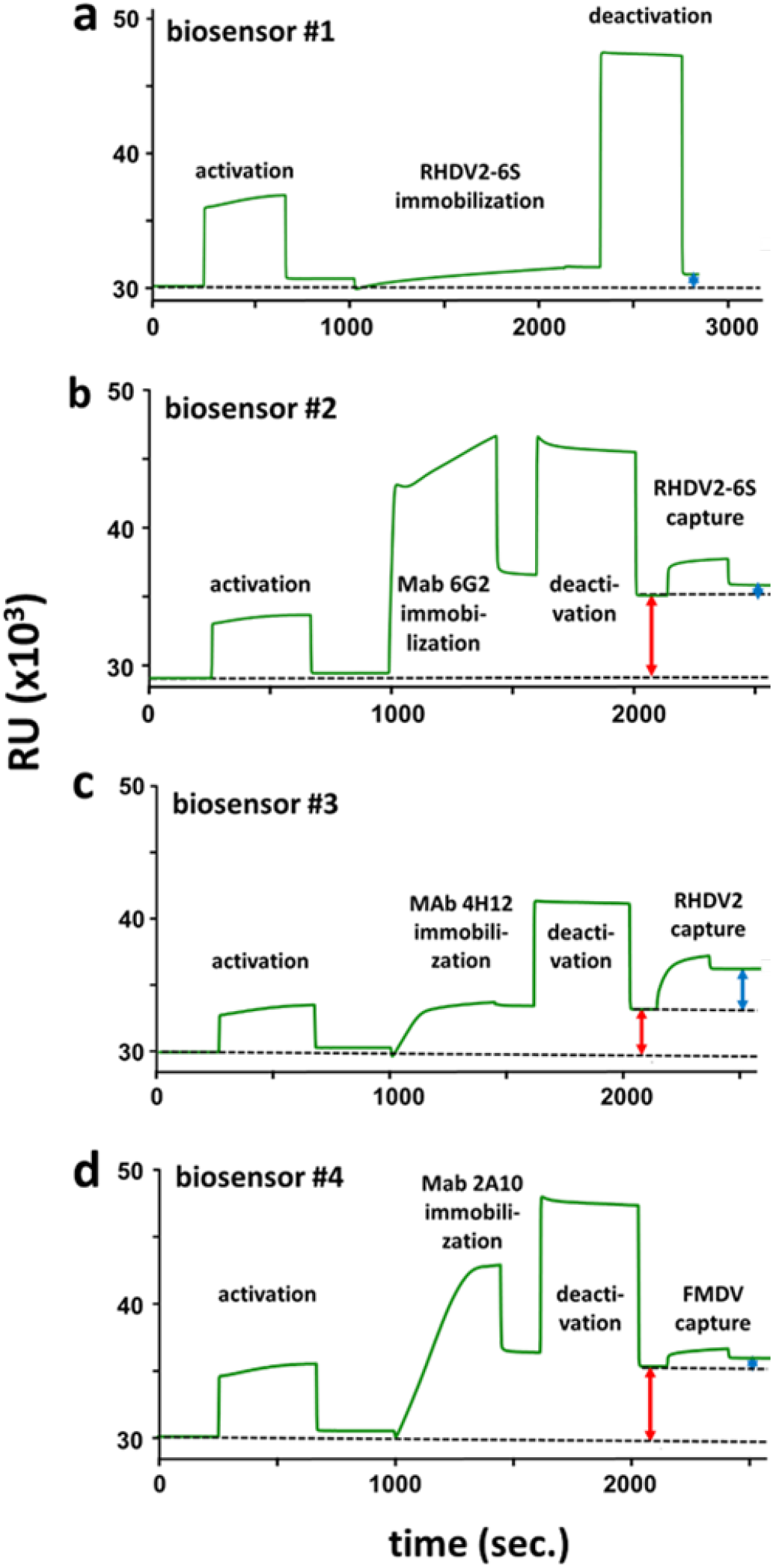
Preparation of the biosensors. Sensorgrams from representative procedures of preparation of biosensor #1, 2, 3 and 4. Dashed line indicates the baseline at the beginning and at the end of the procedures. Red and blue arrows indicate the amount of the catcher-MAb or of the ligand immobilized/captured onto the surface, respectively.

